# A systematic method for detecting abnormal mRNA splicing and assessing its clinical impact in individuals undergoing genetic testing for hereditary cancer syndromes

**DOI:** 10.1101/2022.07.12.499782

**Authors:** Nick Kamps-Hughes, Victoria E.H. Carlton, Laure Fresard, Steve Osazuwa, Elizabeth Starks, John J. Vincent, Sarah Albritton, Robert L. Nussbaum, Keith Nykamp

**Affiliations:** Invitae Corporation, San Francisco, CA, 94103, USA

**Keywords:** RNA sequencing, splicing, variant interpretation, diagnostic testing, splice site prediction

## Abstract

Nearly 14% of disease-causing germline variants result from disruption of mRNA splicing. Most (67%) DNA variants predicted *in silico* to disrupt splicing end up classified as variants of uncertain significance (VUS). We developed and validated an analytic workflow — <u>Sp</u>lice <u>E</u>ffect <u>E</u>vent <u>R</u>esolver (SPEER) — that uses mRNA sequencing to reveal significant deviations in splicing, pinpoints the DNA variants potentially responsible, and measures the deleterious effect of the altered splicing on mRNA transcripts, providing evidence to assess the pathogenicity of the variant. SPEER was used to analyze leukocyte RNA encoding 63 hereditary cancer syndrome genes in 20,317 individuals undergoing clinical genetic testing. Among 3,563 (17.5%) individuals with at least one DNA variant predicted to affect splicing, 971 (4.8%) had altered splicing with a deleterious effect on the transcript and 31 had altered splicing due to a DNA variant located outside our laboratory’s reportable range. Integrating SPEER results into variant interpretation allowed reclassification of VUS to P/LP in 0.4% and to B/LB in 5.9% of the 20,317 patients. SPEER evidence had a significantly higher impact on allowing P/LP and B/LB interpretations in non-White individuals than in non-Hispanic White individuals, illustrating that evidence derived from RNA splicing analysis may reduce ethnic/ancestral disparities in genetic testing.

## INTRODUCTION

Disruption of normal mRNA splicing is a common cause of genetic disease. Approximately 14% of germline pathogenic or likely pathogenic (P/LP) variants cause disease through misplicing^1^, with the vast majority (>90%) of these P/LP splicing variants disrupting the canonical donor and acceptor splice sites (first 1-2 bp flanking an exon) (referred to as CSS). In contrast, many DNA variants other than in the CSS dinucleotides are predicted by commonly used algorithms to affect splicing, but most (69%) of these potential splicing variants (PSpVs) are classified as variants of uncertain significance (VUS)^1^, demonstrating the challenge of classifying non-CSS PSpVs in the absence of confirmatory RNA testing. Knowing which of these PSpVs have a clinically significant impact on splicing could inform variant interpretation, resulting in fewer VUS and increased actionability for genetic testing.

Here, we describe a novel method, the <u>Sp</u>lice <u>E</u>ffect <u>E</u>vent <u>R</u>esolver (SPEER), that uses patient mRNA sequencing to determine whether variants found during DNA sequencing cause aberrant mRNA splicing and have a deleterious impact on transcript structure and function. SPEER accomplishes this task in three steps, first by determining whether there is abnormal splicing compared to controls, then by quantifying by how much normal splicing at a critical splice junction has been reduced, and finally by examining the consequences of abnormal splicing events on mRNA structure and function. SPEER begins with short-read sequencing of cDNA and several overlapping RNA analysis methods^2–6^ to analyze exon-exon junctions and determine whether observed alterations in the mRNA splicing pattern of a gene containing a PSpV are statistically significant relative to a panel of control individuals. In this way, SPEER distinguishes alterations in splicing from the naturally occurring alternative splicing that generates protein diversity and functional specificity across tissue types.^7–9^ Next, it examines the canonical splice junction or junctions most likely impacted by the PSpV and quantifies the extent of loss of normal splicing in the patient compared to normal controls. Finally, SPEER assesses how deleterious the impact of the abnormal splicing of particular mRNA transcripts would be on mRNA stability and function by collecting all the alterations in the transcripts together into an “abnormal splicing event group”, which is then evaluated for reading frame shifts and potential nonsense mediated decay. By assessing the deleterious impact of a DNA variant on transcript structure and function, the evidence generated by SPEER can be used for variant interpretation within a system, such as Sherloc ^10^, that combines many other types of evidence to ultimately assess pathogenicity of a variant.

In this manuscript, we report how we validated SPEER’s ability to detect aberrant splicing events and then applied it to a cohort of more than 20,000 individuals undergoing clinical testing of hereditary cancer syndrome genes to identify which PSpVs detected during DNA sequencing significantly altered splicing and had a deleterious effect on transcript structure and function.

SPEER also detected abnormal splicing changes in a small number of patients, which allowed for the detection and confirmation of DNA variants located outside the reportable range of our next-generation sequencing (NGS) DNA test. When SPEER evidence was used in Sherloc for variant interpretation, a significant fraction of PSpVs could be interpreted definitively as Pathogenic/Likely Pathogenic (P/LP) or Benign/Likely Benign (B/LB); most notably, SPEER allowed an even greater rate of definitive classification of PSpVs among individuals who self-reported as non-White compared to those who self-reported as non-Hispanic White.

## MATERIAL AND METHODS

### Cohorts

In phase 1 of this two-phase study, we ascertained a retrospective cohort of 532 research participants to be used for SPEER validation. Of these 532 participants, 342 had prior genetic testing for hereditary cancer syndromes performed at Invitae and had a germline DNA variant that was known or predicted to alter splicing (PSpV), and 190 did not have a PSpV, but had a personal or family history that was strongly suggestive of a particular hereditary cancer syndrome, e.g. Familial Adenomatous Polyposis (McK #175100). Informed written consent was obtained under a protocol approved by the WCG Institutional Review Board (#20190811). As a reference panel, we used 273 anonymous samples from presumed healthy male and female blood donors (BioIVT, NewYork) that we oversampled for self-reported ancestries underrepresented in genetic studies, including African American (51.2%) and Hispanic (30.4%) individuals. This sampling strategy was designed to guard against mistakenly inferring aberrant splicing when comparing splicing in non-White versus White individals by failing to take into account potential variation in splicing of germline transcripts among individuals of different ancestries, as has been seen in tumor samples in The Cancer Genome Atlas ^11^.

In Phase 2, a prospective cohort of 20,317 individuals referred to Invitae for germline hereditary cancer gene testing underwent paired DNA and RNA sequencing between July 2021 and May 2022. SPEER was used to analyze the leukocyte RNA of 63 hereditary cancer genes in these individuals. Use of de-identified samples and data was approved by an independent institutional review board (WCG IRB #20161796).

### Selection of genes for RNA analysis

A total of 85 hereditary cancer genes were initially considered for RNA analysis (Table S1). Twenty-two of these were ultimately excluded from the final assay because 1) they are expressed at too low a level in leukocyte transcripts to allow statistically significant demonstration of a splicing change, 2) they are genes for which loss-of-function, the usual effect of abnormal splicing, is not a known mechanism for an increased risk for cancer, or 3) they have only a single known loss-of-function missense variant associated with an increased risk for cancer.

### DNA Sequencing

NGS of gene panels were performed as previously described ^1,12,13^ using oligonucleotide baits (Twist Bioscience, South San Francisco, CA; Integrated DNA Technologies, Coralville, IA) to capture coding exon sequences ± 20 bases of flanking intronic sequences, and certain non-coding regions of clinical interest, defined as our reportable range (RR). Targeted regions were sequenced to a minimum depth of 50x and an average depth of 350x read coverage at each nucleotide position within the RR. Full gene sequencing, deletion/duplication analysis, and variant interpretation were performed at Invitae (San Francisco, CA), as previously described ^10,13^.

### Identification of potential splicing variants (PSpV) in DNA

PSpVs were predicted initially using MaxEntScan^14^, SpliceSiteFinder-Like^15^, and the Alamut Splicing Module (Interactive Biosoftware, Rouen, France, version 1.4.4_2016.02.03) and later using SpliceAI ^16^.

### RNA Sequencing

RNA was extracted from whole blood in PaxGene RNA tubes (762165, BD, New Jersey) containing additives that inhibit RNA degradation, including nonsense mediated decay (NMD), using RNAdvance Blood (A35604, Beckman Coulter, Indianapolis) and quantified by fluorometry. Residual DNA contamination was removed by DNaseI treatment (M0303L, New England Biolabs, Massachusetts). Indexed cDNA libraries were prepared using the KAPA hyper RNA kit (KK8541, Roche, Switzerland), substituting proprietary adapters and indexing primers. Transcripts of interest were enriched via hybridization with custom-designed biotinylated oligonucleotide baits (Integrated DNA Technologies, Iowa) followed by Streptavidin bead capture. Illumina compatible P5 P7 sequences were added during post capture amplification and libraries were submitted for 2 × 150bp paired-end next generation sequencing (Illumina, San Diego, CA), targeting 15 million clusters per sample. Adapters were trimmed sequencing reads and UMIs were extracted. Trimmed reads were aligned on the GRCh37 human genome assembly with STAR ^17^, using a RefSeq-based transcript annotation. Duplicate reads were removed using unique molecular identifier sequences (umi-tools,^18^). Raw split read counts were extracted from the deduplicated BAM files using Leafcutter scripts sam2bed.pl and bed2junc.pl ^2^.

### Detection of statistically significant splicing changes

SPEER uses targeted sequencing of RNA isolated from blood samples to determine splice junction counts across 63 genes of interest (Table S1). To assess the statistical significance of splice junction usage in a patient sample, SPEER (Figure 1 A,B) measured usage of known and novel splice junctions using “percent spliced in” (PSI)^19,20^. The PSI of each junction is defined as the ratio of the number of reads supporting the junction divided by the total number of reads overlapping the junction (i.e., supporting reads plus non-supporting reads). SPEER assesses the statistical significance of each splicing change in a patient sample by comparing it to the splice junction usage from the 273-sample reference panel using a beta-binomial test described below.. PSI changes with *p* ≤ 10^−3^ were considered statistically significant. SPEER uses these significant junctions as seeds to recursively find other altered (*p* ≤ 0.05) junctions that share breakpoints in order to link all junctions participating in the localized splicing event into an “abnormal splicing event group” for downstream interpretation.

**Figure 1.**
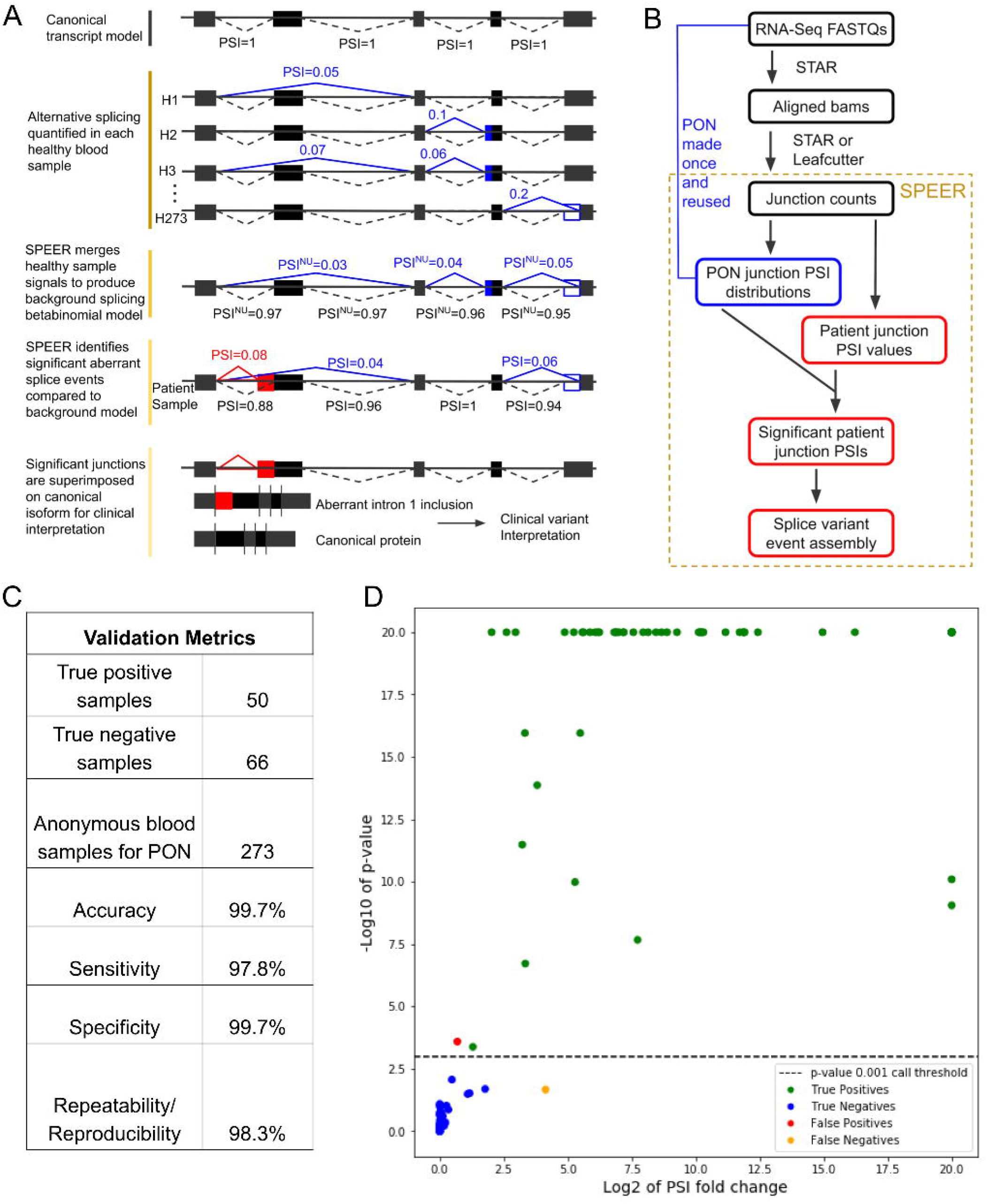
SPEER method and validation. A) SPEER concept. SPEER quantifies the annotated and unannotated splicing variation in the healthy population by modeling the beta-binomial distribution of junction usage for all junctions observed in the healthy normal samples. Patient junction usage is then tested against this distribution to identify junctions that are aberrantly used, filtering out common but unannotated variation. Significant splice aberrations are then mapped to the canonical transcript to infer protein effects. Canonical junctions are shown in dashed black lines, common alternatively-spliced junctions modeled by the normal reference panel are shown in solid blue lines, and the aberrant splice junction detected in the patient sample is shown with a solid red line. B) Bioinformatic workflow. FASTQ files from RNA-Seq experiments are aligned and split reads are extracted and converted to raw junction counts. SPEER uses the raw junction counts from the normal samples to compute the percent spliced in (PSI) distributions for each junction in the healthy normal samples using a beta binomial count model. Beta binomial parameters for each junction are saved in a panel of normal splicing (PONS) file, which is calculated once and reused for every process-matched patient sample. Patient junction counts are converted to PSI values, which are tested against the panel of normal samples to identify significantly aberrant junction usage. Significant patient junctions are positionally linked to assemble splice variant events. C) Validation metrics. SPEER was validated using a set of positive and negative control blood samples from patients with known and expected splicing events. Anonymous blood samples were used to create the PONS file. SPEER showed high accuracy and reproducibility in detecting abnormal splicing events in patient samples. D) Quantitative features of abnormal splicing events used for validation. SPEER p-value and PSI fold change versus PONS are plotted for known positive and negative control splice junctions. Positive and negative calls are strongly separated and inaccurate calls fall near the p-value threshold. Infinite values are set to 20 on both axes for display purposes and the top right point contains 11 samples with infinite fold change and a p-value of 0.

### Beta-binomial model description and parameter estimation

For every junction observed in at least one control, a PSI normal model was calculated using the beta-binomial distribution. In this model, the number of reads overlapping the junction was considered the number of trials and the reads supporting the junction was considered the number of successes. For each junction, the parameters nu (approximately the mean PSI) and rho (a measure of the increased variance relative to the binomial distribution) were fit to their maximum likelihood values and stored as a panel of normal splicing bed file. The control parameters for each junction were then compared to the corresponding PSI for each patient sample to generate a p-value using a beta-binomial likelihood ratio test. This statistical approach was inspired by the ShearwaterML algorithm for detecting low frequency somatic DNA variants from NGS data ^21^.

### Quantifying the loss of the normal transcript directly impacted by a PSpV

Each abnormal splicing event group observed in a patient sample included a reduction in the PSI for one or more canonical splice junction and an increase in PSI for one or more newly created or non-canonical splice junction (see Figure 1A). To measure the reduction in normal splicing associated with a PSpV, we calculated the ratio of the PSI for the canonical junction determined to be directly impacted by the PSpV over the PSI for that same junction in the control panel. This metric, which we define as PSI-X, is expected to correlate with loss of the normal or most commonly expressed transcript. Moreover, a significant reduction in PSI-X is expected to correlate with pathogenicity for PSpVs in genes known to cause disease with an autosomal dominant, loss-of-function mechanism.

Directly impacted canonical junctions were defined as the following: for intronic PSpVs, we chose the canonical junction that was normally created by splicing out the intron that contained the PSpV; for exonic PSpVs we chose the canonical junction with the highest PSI in the control panel that also shared a breakpoint with a newly created, non-canonical junction. If the highest PSI value for a canonical junction was less than 2x other canonical junctions with shared breakpoints, the PSI values were averaged across all impacted canonical junctions.

### Assessing whether abnormal splicing events impact translation and function of the protein

Abnormal splicing event groups with a reduction in PSI-X that had at least one breakpoint within 50 bp of a PSpV were flagged for review by trained variant scientists (Supplemental Methods). This was an essential step for assessing whether the transcript created by non-canonical junctions may compensate for the reduction in normally spliced mRNA. As an example, a PSpV that abolishes a well annotated acceptor splice may result in substantially decreased PSI-X while also activating a cryptic acceptor splice site three nucleotides upstream, resulting in a novel protein with a single amino acid insertion. It’s uncertain whether this novel protein will compensate for the loss of the normally expressed protein, thus warranting a VUS classification for the PSpV. The first step in this review was to label each splice junction in an event group with one of the following tags: (1) *canonical*, a junction formed by adjacent exons in the full-length canonical transcript); (2) *exon skipping*, defined as non-adjacent junctions with one more exons being skipped; (3) *partial exon exclusion*, in which a portion of a canonical exon was not included in the transcript; (4) *partial intron inclusion*, in which a portion of an intron was included in a transcript; (5) *cryptic exon*, when two non-canonical junctions flanked an intronic sequence. All event groups, other than complex, were labeled according to whether the primary non-canonical junction is expected to cause nonsense-mediated decay (NMD). Event groups with a single non-canonical junction causing a frameshift and/or premature termination codon upstream of the last coding exon were labeled as *NMD+*. Those expected to result in in-frame insertions anywhere in the transcript, or in deletions or premature termination codons downstream of the last exon junction, were labeled as *NMD-*. If an event group had multiple non-canonical junctions, with different predicted effects on protein translation, the event group was labeled as *complex*.

### Detection and classification of variants outside our test’s reportable range

Rare variants located outside our test’s reportable range (all coding exons plus 20 bp flanking each exon) are missed by standard panel-based NGS tests. When SPEER identifies an abnormal splicing event group at the variant discovery significance threshold (i.e., *p* ≤ 10e^-5^) *and* one or more abnormal junctions exhibit a fold-change in PSI ≥10,000, we manually examine the primary NGS data for a DNA variant that may explain the abnormal splicing event. If none is found in the primary data, we perform long-read DNA sequencing (PacBio, CA) throughout the region surrounding the abnormal splicing event. If a PSpV is identified in the region surrounding the abnormal splicing event, it is flagged for interpretation by the variant scientist team. If a PSpV could not be identified, the abnormal splicing change was presumed to be a technical false positive or the result of natural variation in splicing and not included in the clinical report.

### Integrating deleterious RNA splicing changes as evidence in DNA variant classification

In order to standardize the application of SPEER evidence for variant classification, we created a new category of evidence within our semi-quantitative variant classification framework, Sherloc^10^. We referred to this evidence category as *Observed RNA Effects* (see Table S2).

Each criterion within this new evidence category was assigned pathogenic (P) or benign (B) point scores. Point scores were assigned and calibrated by applying Observed RNA Effects criteria to more than 300 PSpVs variants observed in the retrospective cohort of 532 research participants in Phase 1. Final classifications in Phase 2 of the study were assigned based on the sum of all available evidence, including the Observed RNA Effects criterion when applicable.

Rates of PSpV upgrades (VUS to LP/P) and downgrades (VUS to LB/B) observed during phase 2 of the study were calculated by removing Observed RNA effect criteria and asking whether the final classification (and Sherloc score) moved from P/LP or B/LB to a VUS. Differences in these classification rates among self-reported ethnicities were tested using a one-sided Fisher Exact Test by comparing the proportion of individuals with downgrades and upgrades for each ethnicity (Black/African-American, Hispanic, Asian and Other) to the proportion of individuals from the most common self-reported ethnicity (non-Hispanic White).

## RESULTS

### SPEER validation

SPEER was validated using 50 positive control samples containing 46 unique DNA variants as follows: 10 samples with 7 unique variants well documented to alter splicing according to published literature; 13 samples with 12 unique variants in the essential GT or AG dinucleotides but without published functional evidence; and 27 samples with 27 unique variants located within 50 nucleotides of a significant abnormal splicing event (*p* ≤ 0.001). False positives were any significant abnormal splicing event (*p* ≤ 0.001) detected throughout the 63-gene panel for a canonical junction not within 50 bp of a known PSpV. True negatives (TN) were derived from two sets of samples. The first set were the same 50 samples that had served as *true positives* for one junction-variant pair, repurposed as a source of *true negatives* by analyzing all other splice junctions lacking a nearby PSpV. The second set of true negative samples consisted of 16 samples with a PSpV that nonetheless yielded a normal splicing pattern based on a manual review of RNA sequence alignments to establish the absence of a splicing change. Combining the two sets yielded 66 true negative (TN) samples suitable for calculating sensitivity, specificity, accuracy, reproducibility of SPEER’s findings between replicates of the same sample analyzed in parallel and in the same sample over time (Figure 1 C,D and Table S3).

SPEER detected a significant splicing change in 45 of 46 positive control DNA variants in the 50 positive control samples (97.8% sensitivity; Figure 1C) while maintaining a specificity of 99.7%. We used two of the remaining four positive control samples, along with eight of the negative control samples, to perform reproducibility and repeatability testing in triplicate across all 10 samples. All 30 (100%) of the reproducibility samples and 29 (96.7%) of 30 repeatability samples yielded the expected results (Figure 1C). True positive and true negative calls demonstrated strong separation at the p-value threshold used for detecting splicing effects (Figure 1D), and the same variants found in multiple samples demonstrated consistent PSI values (data not shown). SPEER also enabled the detection of variants outside of the reportable range in four individuals from the phase 1 research cohort, all of whom had presented with a persuasive personal and/or family history of hereditary cancer syndrome.

### Characterization of DNA variants associated with abnormal splicing event groups

SPEER detected abnormal splicing events associated with PSpVs with very high accuracy (99.7%). The next step was to calculate PSI-X as a measure of the loss of normal splicing at the canonical junction most relevant to the PSpV compared to controls. We plotted the distribution of PSI-X for variants previously classified as P/LP due to a splicing abnormality, and therefore could be confidently considered to have a deleterious effect on the transcript, and those known to be B/LB and therefore could be considered to have no deleterious effect on splicing (Figure 2A). We computed a receiver operating characteristic (ROC) curve (Figure 2B) for these data that resulted in an area under the ROC curve (AUROC) of 0.91 for PSI-X, demonstrating that PSI-X has excellent discriminatory power for separating PSpVs known to be either deleterious or not, inferred from their status as either P/LP or B/LB.

**Figure 2.**
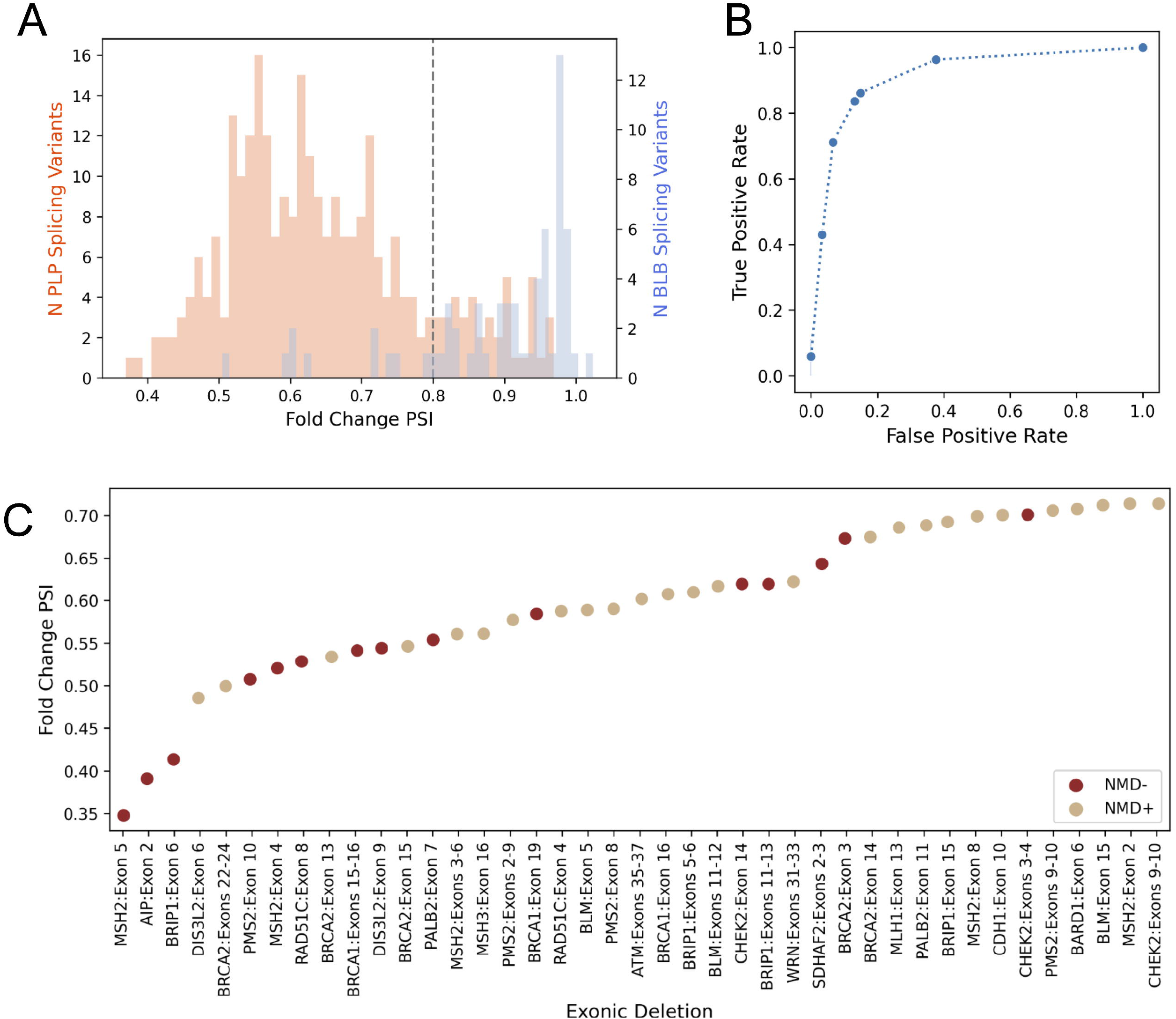
Establishing a PSI-X threshold for interpreting abnormal splice events as pathogenic. A) PSI-X (percent spliced in fold change) distribution for PSpVs (potential splicing variants) classified as pathogenic or likely pathogenic (PLP, beige) or benign or likely benign (BLB, blue). Dotted line represents PSI-X of 0.80. B) Receiver operating characteristic (ROC) curve for the data in panel A. The area under the ROC curve is 0.91. C) PSI-X values for heterozygous exon deletions detected by DNA sequencing. Although the PSI-X for deletions expected to cause NMD (nonsense mediated decay, beige circles, NMD+) is statistically different (one-sided t-test, p=0.0026) than the PSI-X for deletions expected to escape NMD (red circles, NMD-), all events are below 0.8, demonstrating the ability to correctly identify them independently of NMD status.

As another line of evidence to validate the PSI-X metric for quantifying the effect of a variant on reducing splicing of the junction most relevant to that variant, we assessed PSI-X for samples in which a heterozygous DNA deletion encompassing one or more exons that would therefore represented complete loss of junction usage for the flanking splice sites. With these deletion variants we found a distribution in PSI-X from 0.3 to 0.75 (Figure 2C), suggesting that a PSI-X <0.8 was strongly correlated with significant reduction in the usage of a canonical splice junction from one allele.

We then sought to assess how well PSI-X alone could predict what would be the ultimate clinical interpretation of a PSpV as either P/LP or B/LB. If all PSpVs with a PSI-X <0.8 are deleterious to mRNA transcript structure and function and therefore might be naively assumed to be P/LP, and all PSpVs with a PSI-X ≿0.8 were considered non-deleterious and therefore taken to be B/LB, a comparison of predicted interpretation based on PSI-X with the known pathogenicity status of these variants would be 100% accurate. This is not the case as PSI-X alone had an accuracy of 86% for the clinical interpretation of variants. Clearly, not all deleterious changes in mRNA structure and function must be pathogenic and not all variants that leave splicing intact are necessarily benign. We therefore included the likelihood that an abnormal splicing event group would lead to NMD, thereby affecting mRNA stability and expression of the translated protein product, in the analysis of PSpVs with PSI-X <0.8. In this way, we recognized that six of nine known B/LB variants with a PSI-X <0.8 affect transcript structure but are unlikely to affect gene function, since the abnormal transcripts created were not expected to result in NMD; five event groups resulted in small in-frame (≤ 6 amino acids) deletions or insertions, while one event group resulted in alternative usage of the first coding exon. By combining a requirement that the PSI-X be <0.8 *with* the requirement that the abnormal transcripts lead to NMD, we could achieve a positive predictive value (PPV) for PSI-X <0.8 of 98.9% for P/LP variants, sufficiently high to be used as evidence for pathogenicity in Sherloc. In contrast, among 42 PSpVs with PSI-X > 0.8, 9 variants were previously known to be P/LP presumably because they disrupted the protein directly through coding sequence changes (i.e., nonsense, missense, in-frame deletion or insertion) and not through abnormal splicing. Therefore, by combining a PSI-X ≿0.8 with the requirement that the PSpV not be a missense variant or directly change the coding sequence, we could improve the negative predictive value (NPV) of PSI-X > 0.8 for the variant being B/LB to 61.2%, which we did not consider sufficiently high to be used as evidence for the variant being benign in the Sherloc classification framework. More work needs to be done to increase the specificity of PSI-X ≿0.8 for B/LB variants.

### Prospective analysis of paired DNA and RNA testing

In Phase 2 of the study, 20,317 individual blood samples were processed for DNA sequencing and for RNA extraction and sequencing. Across the entire population, 3,563 (17.5%) of the patients had at least one DNA variant identified by NGS that was a PSpV (Table 1). Abnormal splicing, defined by a statistically significant event group, coincided with 605 unique PSpVs (Table 2, Row 1) in a total of 971 patients (4.8% of all prospective samples) (Table 1). These 605 PSpVs were detected in 54 of the 63 genes analyzed by SPEER (Table S4). We observed normal splicing associated with a PSpV in 2,487 patients (73% of patients with a PSpV), illustrating the high false positive rate for the algorithms and heuristics used to identify PSpVs.

**Table 1:**
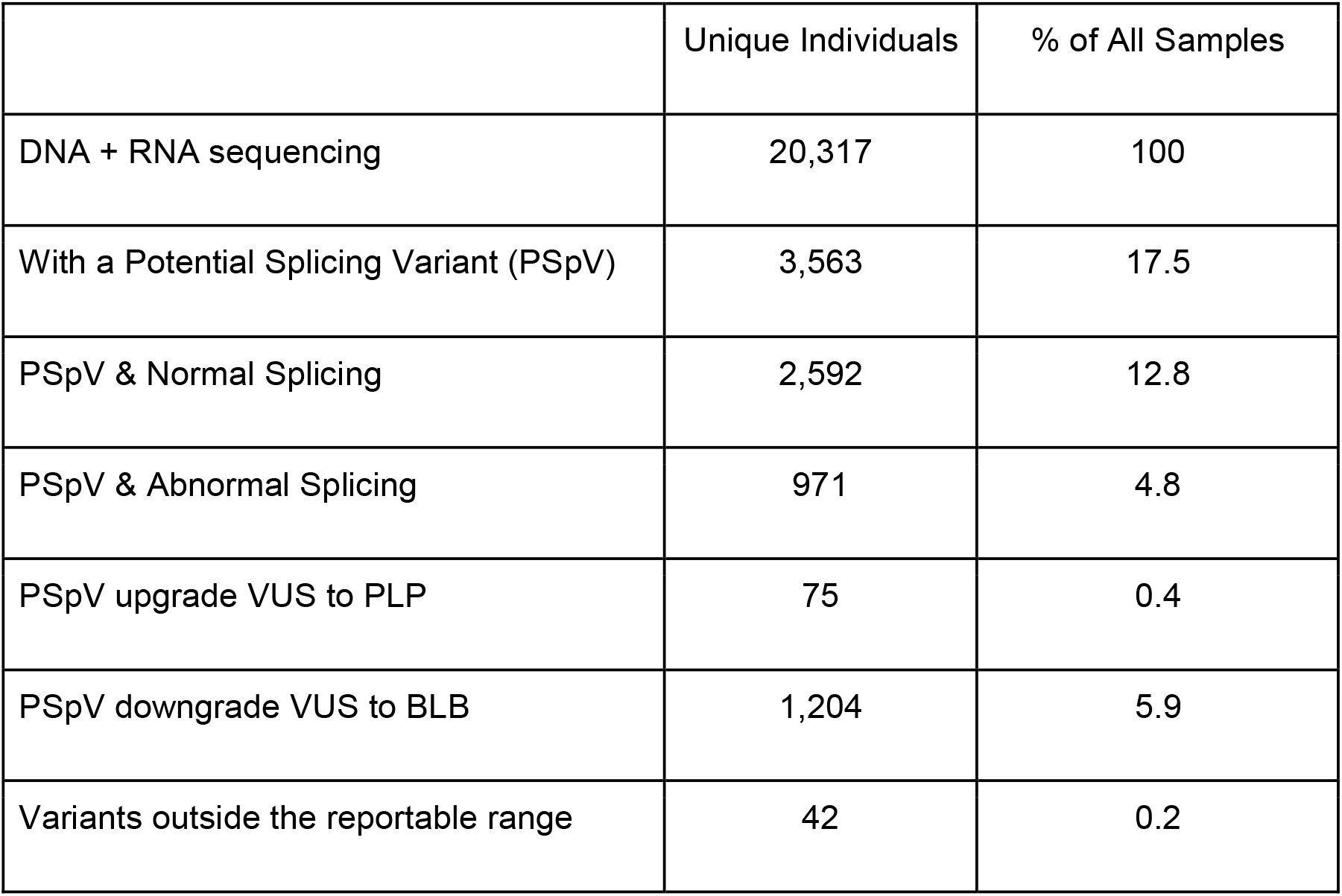
Results from Phase 2.

|  | Unique Individuals | % of All Samples |
| --- | --- | --- |
| DNA + RNA sequencing | 20,317 | 100 |
| With a Potential Splicing Variant (PSpV) | 3,563 | 17.5 |
| PSpV & Normal Splicing | 2,592 | 12.8 |
| PSpV & Abnormal Splicing | 971 | 4.8 |
| PSpV upgrade VUS to PLP | 75 | 0.4 |
| PSpV downgrade VUS to BLB | 1,204 | 5.9 |
| Variants outside the reportable range | 42 | 0.2 |

**Table 2:** Classification of PSpVs in the Phase 2 study.

|  | PLP | VUS | BLB | Total PSpVs |
| --- | --- | --- | --- | --- |
| PSpV & Abnormal splicing | 260 (42.9%) | 275 (45.5%) | 70 (11.6%) | 605 (100%) |
| PSpV & Normal splicing | 120 (9.4%) | 541 (42.3%) | 617 (48.3%) | 1278 (100%) |
| Total PSpVs | 380 (20.2%) | 816 (43.3%) | 687 (36.5%) | 1883 (100%) |

By evaluating the location of PSpVs in relation to the exonic sequence, we found that the largest number of P/LP variants relative to VUS were within the CSS (−1, -2, +1, +2; Figure 3A). In addition, there was an enrichment at positions +3, +4, and +5 from the canonical donor splice site, and a cluster at -11, -10 and -9, upstream of the canonical acceptor site that most often creates a novel acceptor splice site. Overall, we observed abnormal splicing events associated with 605 PSpVs (Table 2), with 93% of these PSpVs located within our test’s reportable range of +/- 20 bp flanking the exons.

**Figure 3.**
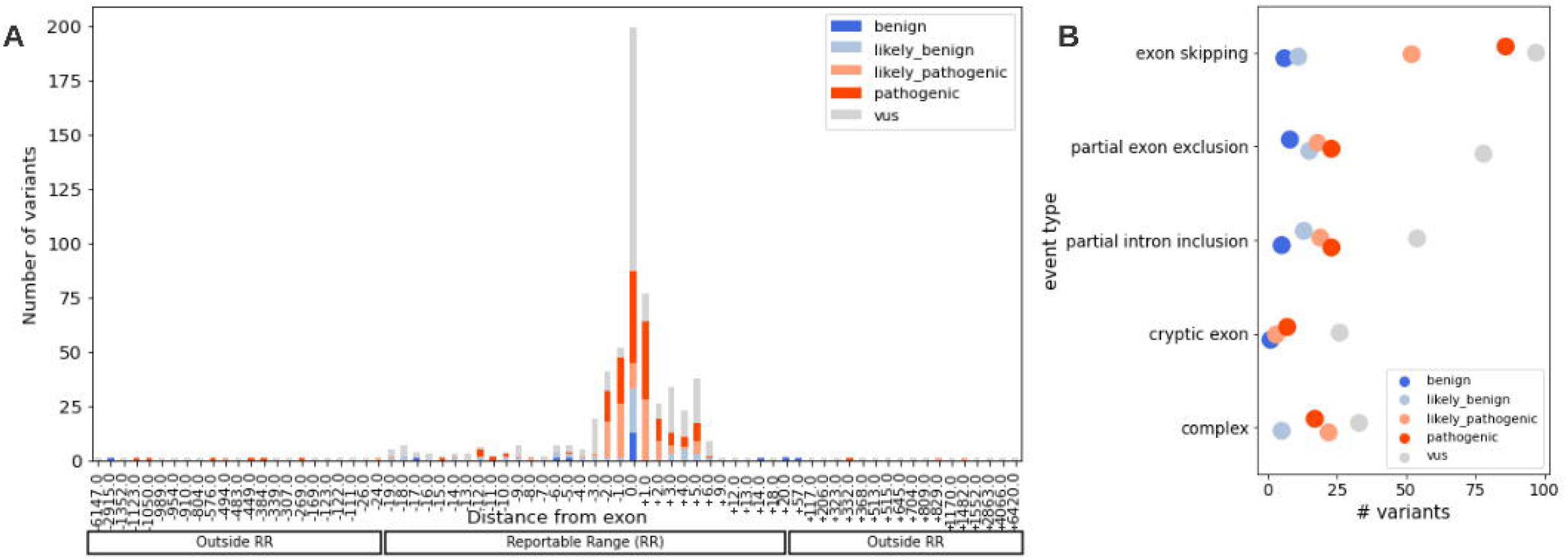
Characterization of splicing events observed in the prospective cohort. A)Position of variants associated with splicing events in regards to the transcript exon (0 is exonic). Although there is a clear enrichment of likely pathogenic and pathogenic variants at +1, +2 and +3, we do observe both benign and pathogenic variants at other positions. Overall, 93% of PSpVs associated with abnormal splicing events are within 20 bps of the exon, corresponding with the reportable range of the NGS DNA test. B) Splicing event types plotted by the number of potential splicing variants (PSpVs) observed. Colors represent classification of PSpVs following interpretation of abnormal splicing events. Most PSpVs classified as pathogenic or likely pathogenic are associated with changes leading to exon-skipping, while PSpVs associated with partial exon exclusion, partial intron inclusion, and complex events show the largest discrepancy between definitive classifications and variants of uncertain significance.

### Classification of abnormal splicing events in the prospective cohort

Each PSpV associated with an abnormal splicing event group identified by SPEER was classified using Observed RNA Effects evidence within the Sherloc interpretation framework. Overall, half (56.7%) dof predicted splicing variants (PSpVs) observed in this study have been classified as pathogenic or likely pathogenic (P/LP; 20.2%) and benign or likely benign (B/LB; 36.5%), while the remaining (43.3%) were classified as variants of uncertain significance (VUS) (Table 2, row 3). As expected, abnormal splicing correlated with a much higher rate of P/LP variants (42.9%) and much lower rate of B/LB variants (11.6%) (Table 2, row 1). Conversely, normal splicing correlated with a much lower rate of P/LP variants (9.4%) and much higher rate of B/LB variants (48.3%) (Table 2, row 2). Importantly, we found that 6.3% of all patients from the phase 2 cohort had a P/LP or B/LB variant that would have been a VUS by our interpretation framework if it were not for the added RNA splicing data from our study: 75 patients (0.4% of all samples) would have received a VUS instead of a P/LP classification, and 1,204 patients (5.9% of all samples) would have received a VUS instead of a B/LB classification (Table 1). When comparing these data across self-reported ancestries, we found that individuals with self-reported Hispanic (9.4%, p=8.2e-12), Black/African American (10.4%, p=1.8e-16), or Asian (9.1%, p=1.4e-5) ancestries had statistically significantly higher rates of definitive classifications (P/LP or B/LB) than Non-Hispanic White populations (4.8%) due to the addition of supplemental RNA splicing data (Table 3).

**Table 3:** Reclassification rates among self-reported ethnic populations.

|  | Unique individuals | % of ethnic population |
| --- | --- | --- |
| Non-Hispanic White | 608 | 4.8 |
| Black/African American | 158 | 10.4* |
| Asian | 55 | 9.12* |
| Hispanic | 137 | 9.4* |
| Other** | 312 | 9.7 |

\*p-value<0.05 after one-sided Fisher Exact test. \*\*Includes Ashkenazi Jewish, Native American,
Multiple and Unknown.

The most abundant splicing change in the prospective cohort was exon skipping, with partial exon exclusion and partial intron inclusion events being the next most abundant events. Partial exon exclusion, partial intron inclusion, and complex events displayed the largest discrepancies between PSpV definitive classifications (i.e., P, LP, LB, B) and VUS, indicating that these were the most challenging event groups to interpret (Figure 3B).

### Detection of variants affecting splicing outside the reportable range

Among the 190 individuals ascertained in the initial research cohort in Phase 1 specifically because no P/LP variants were found through routine clinical testing despite a personal or family history highly suggestive of a hereditary cancer syndrome, 4 (2.1%) were found to have a variant outside our reportable range with a deleterious effect on splicing. In the prospective consecutive cohort of 20,317 patients referred for hereditary cancer testing, 42 (0.2%) patients carried 35 unique variants outside of the reportable range that could be associated with abnormal splicing (Table 1). Of these 35 variants, 6 (17.1%), seen in nine patients, were classified as P/LP, representing a 0.04% yield of P/LP variants not present in the reportable range of our test in the entire cohort (9 of 20,317 patients). An additional two variants, seen in two patients (4.8%), were classified as benign, and 27 of the 35 unique variants, present in 31 patients (73.8%), were classified as VUS (Figure 4).

**Figure 4.**
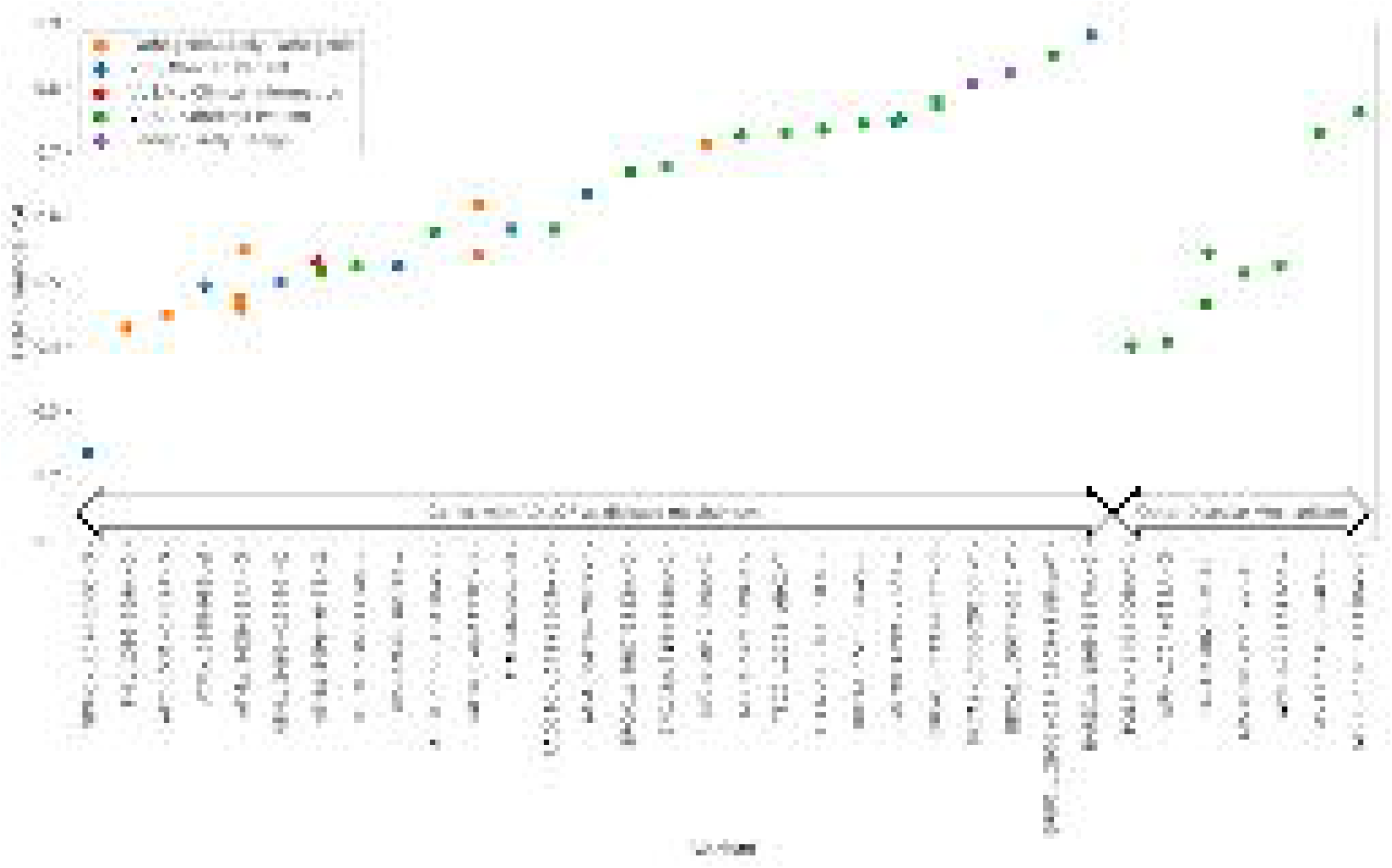
Discovery of intronic variants outside our reportable range with SPEER. Thirty-five intronic DNA variants outside our reportable range were identified in this study. Five DNA variants were classified as pathogenic or likely pathogenic (orange), two DNA variants were classified as benign or likely benign (purple), and the remaining were classified as variants of uncertain significance (VUS). The affected status of individuals with VUS is represented by color: green = unaffected, blue = affected, and red = unknown due to lack of clinical information provided. The canonical PSI fold change (PSI-X) showed a large distribution for these abnormal splicing events, ranging from 0.23 to 0.9.

## DISCUSSION

We describe SPEER, an accurate and scalable approach for detecting and quantifying splicing alterations in peripheral blood leukocyte transcripts of 63 hereditary cancer syndrome genes in patients undergoing genetic testing for these syndromes. We demonstrate that adding RNA analysis to DNA sequencing of hereditary cancer syndrome genes facilitates the interpretation of DNA variants expected or predicted to alter splicing. In addition, of the 605 unique variants identified in the prospective cohort that affected splicing, 35 of them (5.8%) were located outside the reportable range of +/- 20 bp flanking the exons of our NGS panel tests and were detected by SPEER.

We found that supplemental RNA analysis led to a definitive classification of DNA variants that would have been classified as a VUS without the RNA analysis in 6.3% of individuals in a prospective cohort undergoing testing for hereditary cancer; of these, 0.4% received a P/LP classification while 5.9% of patients tested had a B/LB variant, thereby effectively reducing VUS rates. This fraction of patients with reclassifications due to RNA analysis in our 20,317 prospective cohort was higher (6.3%) than what was recently reported in a cohort of 43,524 patients undergoing DNA and RNA testing for hereditary cancer (1.3%) ^22^. This difference is likely because of the larger number of genes (63 genes) tested by RNA in our cohort compared to the recently published cohort (18 genes). Importantly, the rate of definitive classifications (i.e., P, LP, LB, B) dependent on RNA analysis was significantly higher in Black/African American, Asian, and Hispanic individuals combined than in non-Hispanic White individuals. The preferential impact of RNA analysis on VUS reclassification in these less-studied populations probably resulted from RNA analysis providing direct evidence that was not limited by the insufficient representation of individuals of non-European origin in variant databases or in the published literature. This indicates that including RNA analysis in the interpretation of DNA variants predicted to impact splicing may preferentially reduce VUS rate for less-studied ancestral and ethnic populations, thereby reducing health disparities.

These results also highlight the nuances and difficulties of incorporating sample-based RNA splicing analysis into variant interpretation frameworks used in routine genetic testing. Alternative splicing is very common, and it can be challenging to pinpoint specific splicing alterations associated with a hereditary disease. For example, greater than 96% of all patient samples in our prospective cohort had at least one statistically significant splicing change at the p ≤ 0.001 threshold compared to the reference panel of normal samples. This statistical threshold was not multiple hypothesis-corrected because it was meant to accurately determine whether a specific PSpV altered normal splicing. By focusing scientific review on splicing alterations with a PSpV within 50 bp of the nearest altered junction, we reduced the frequency of abnormal splicing event groups to 4.8% of all patient samples, while maintaining 99.7% accuracy for detecting abnormal splicing events previously known to be associated with DNA variants in our validation of SPEER.

Even with supplemental RNA splicing data, the biological impact and clinical relevance of DNA variants associated with a significant splicing alteration are difficult to assess, often still resulting in VUS classifications. This is particularly true for variants found outside of the reportable range of the test since they are often observed in a single individual and lack the additional corroborating evidence needed to make a definitive classification, such as population frequency data in publicly available databases. Only 9 of 20,317 patients (0.04%) in our prospective cohort had variants outside our +/-20 bp reportable range, caused altered splicing, and would ultimately be classified as P/LP. This rate of P/LP variants that occur >20bp into the introns is consistent with the rate (0.06% or 28/43,524) seen in another large RNA cohort undergoing testing for hereditary cancer ^22^. Another 2 of 20,317 patients had variants outside our +/-20 bp reportable range, caused altered splicing, but still were ultimately classified as B/LB, underscoring the fact that a variant that has a deleterious effect on splicing may still not cause disease.

Another complicating factor in connecting a deleterious effect on splicing to pathogenicity is that abnormal splicing can be incomplete. A heterozygous DNA variant may result in a reduction of normal mRNA splicing that ranges from 0% (i.e., no reduction) to 50% (i.e., complete loss of normal splicing from the variant allele), with everything in between considered partial loss, also referred to as ‘leakiness^23,24^.Importantly, because a PSpV that leads to a complete loss of normal splicing is more likely to cause disease than a PSpV that leads to a partial loss, accurate classification of PSpVs using abnormal splicing data requires a confident measure of the amount of reduction. We developed such a measurement to quantify the reduction in normal mRNA splicing — PSI-X — and determined that splicing alterations in blood samples associated with a PSI-X <0.8 correlate with sufficient loss of normal splicing from one allele to cause disease and qualify as pathogenic evidence for autosomal dominant hereditary cancer syndromes within our variant classification framework. Although a PSI-X threshold of 0.8 demonstrates high discriminatory performance (AUROC = 0.91) for PSpVs with abnormal splicing events, we still find that 15% of B/LB variants fall below the threshold and 14% of P/LP variants are above the threshold. To achieve a level of confidence in the final classification of PSpVs associated with abnormal splicing suitable for clinical testing, it is essential that trained variant scientists review the expected effect of abnormal splicing on translation and protein function, and then integrate these findings into a DNA variant interpretation framework. In this way, we demonstrated that by pairing DNA and RNA testing, we can increase the number of patients who have an actionable finding while achieving a PPV of 98.9%. On the other hand, even with a careful review of the abnormal event groups associated with PSI-X ≿0.8, it was still difficult to discriminate between P/LP and B/LB variants, as demonstrated by a much lower NPV (61.2%) when PSI-X is≿0.8. As a result, to limit the likelihood of a false negative result for our clinical testing patients, we chose not to award benign points in the Sherloc framework even for statistically significant abnormal splicing event groups when PSI-X ≿0.8. PSpVs associated with these higher PSI-X values will generally be VUS unless complementary evidence (i.e., family segregation, population frequency, additional splicing studies) support a more definitive classification.

Although this study clearly demonstrates that RNA splicing data can aid in variant classification and identification, the results should be interpreted in the context of the following technical and biological limitations. First, the 63 hereditary cancer genes included in this study are tumor suppressor genes that are not highly expressed in whole blood leukocyte RNA. We expect each gene to have tissue-specific expression and alternatively spliced products that differ from what can be detected in a blood sample. As a result, abnormal splicing observed from whole blood leukocytes may not reflect a biological change in the tissue of interest, while the absence of a splicing alteration in whole blood may not always correlate with an absence of abnormal splicing in the disease-relevant tissue. Second, the SPEER algorithm is designed to evaluate changes in junction usage. A small fraction of splicing alterations result in complete intron retention, which escapes detection by the SPEER algorithm in its current form. This likely explains why, in our study, several known P/LP DNA variants had PSI-X ≿0.8 with transcripts that underwent NMD. Further development is underway to include a method for evaluating the relative frequency of complete intron inclusion in the SPEER algorithm, which will better resolve the uncertainty of variants with high PSI-X. Third, although PSI values and aberrant junctions were highly consistent for the same variant across multiple samples, a few DNA variants showed high levels of variability. Most of these correlated with the presence of *cis*-acting elements, such as secondary DNA variants, suspected of modifying the impact of the primary DNA variant on splicing. Currently it is not possible to predict *a priori* which splicing changes are impacted by modifiers, but with more samples sequenced for the same DNA variants we may be able to identify patterns in the DNA sequence that modulate the PSI and alternative splicing events within these tumor suppressor genes.

In summary, RNA analysis in whole blood is a valuable tool for finding and assessing the impact of DNA variants suspected of affecting RNA splicing, thereby altering gene expression. It helps to reduce VUS rates in a fraction of individuals who have DNA variants predicted to affect splicing, and supports the identification of deleterious, possibly pathogenic variants outside the test’s reportable range. However, interpretation of the splicing information is challenging and there are limitations in the techniques and in our biological understanding of splicing. By combining comprehensive DNA and RNA sequencing with the validated SPEER algorithm, we will continue to collect data individuals referred for hereditary cancer syndrome testing, develop deeper insights into the clinical relevance of splicing alterations, and improve the interpretation of inherited DNA variants — leading to a further reduction in uncertainty for hereditary cancer gene testing.

## Notes

### Competing Interest Statement

All authors are employed by and shareholders of Invitae, Inc.

### Summary of Updates

The manuscript was revised for clarity and important methods were moved to the primary body of the text.

## REFERENCES

1. Truty R, Ouyang K, Rojahn S, Garcia S, Colavin A, Hamlington B, Freivogel M, Nussbaum RL, Nykamp K, Aradhya S. Spectrum of splicing variants in disease genes and the ability of RNA analysis to reduce uncertainty in clinical interpretation. Am J Hum Genet, 2021, 108:696–708

2. Li YI, Knowles DA, Humphrey J, Barbeira AN, Dickinson SP, Im HK, Pritchard JK. Annotation-free quantification of RNA splicing using LeafCutter. Nat Genet, 2018, 50:151–8

3. Frésard L, Smail C, Ferraro NM, Teran NA, Li X, Smith KS, Bonner D, Kernohan KD, Marwaha S, Zappala Z, Balliu B, Davis JR, Liu B, Prybol CJ, Kohler JN, Zastrow DB, Reuter CM, Fisk DG, Grove ME, Davidson JM, Hartley T, Joshi R, Strober BJ, Utiramerur S, Lind L, Ingelsson E, Battle A, Bejerano G, Bernstein JA, Ashley EA, Boycott KM, Merker JD, Wheeler MT, Montgomery SB. Identification of rare-disease genes using blood transcriptome sequencing and large control cohorts. Nat Med, 2019, 25:911–9

4. Mertes C, Scheller IF, Yépez VA, Çelik MH, Liang Y, Kremer LS, Gusic M, Prokisch H, Gagneur J. Detection of aberrant splicing events in RNA-seq data using FRASER. Nat Commun, 2021, 12:529

5. Jenkinson G, Li YI, Basu S, Cousin MA, Oliver GR, Klee EW. LeafCutterMD: an algorithm for outlier splicing detection in rare diseases. Bioinformatics, 2020, 36:4609–15

6. Vaquero-Garcia J, Barrera A, Gazzara MR, González-Vallinas J, Lahens NF, Hogenesch JB, Lynch KW, Barash Y. A new view of transcriptome complexity and regulation through the lens of local splicing variations. Elife, 2016, 5:e11752

7. Sibley CR, Blazquez L, Ule J. Lessons from non-canonical splicing. Nat Rev Genet, 2016, 17:407–21

8. Baralle FE, Giudice J. Alternative splicing as a regulator of development and tissue identity. Nat Rev Mol Cell Biol, 2017, 18:437–51

9. Ule J, Blencowe BJ. Alternative Splicing Regulatory Networks: Functions, Mechanisms, and Evolution. Mol Cell, 2019, 76:329–45

10. Nykamp K, Anderson M, Powers M, Garcia J, Herrera B, Ho Y-Y, Kobayashi Y, Patil N, Thusberg J, Westbrook M, Invitae Clinical Genomics Group, Topper S. Sherloc: a comprehensive refinement of the ACMG-AMP variant classification criteria. Genet Med, 2017, 19:1105–17

11. Al Abo M, Hyslop T, Qin X, Owzar K, George DJ, Patierno SR, Freedman JA. Differential alternative RNA splicing and transcription events between tumors from African American and White patients in The Cancer Genome Atlas. Genomics, 2021, 113:1234–46

12. Truty R, Paul J, Kennemer M, Lincoln SE, Olivares E, Nussbaum RL, Aradhya S. Prevalence and properties of intragenic copy-number variation in Mendelian disease genes. Genet Med, 2019, 21:114–23

13. Kurian AW, Hare EE, Mills MA, Kingham KE, McPherson L, Whittemore AS, McGuire V, Ladabaum U, Kobayashi Y, Lincoln SE, Cargill M, Ford JM. Clinical evaluation of a multiple-gene sequencing panel for hereditary cancer risk assessment. J Clin Oncol, 2014, 32:2001–9

14. Yeo G, Burge CB. Maximum entropy modeling of short sequence motifs with applications to RNA splicing signals. J Comput Biol, 2004, 11:377–94

15. Shapiro MB, Senapathy P. RNA splice junctions of different classes of eukaryotes: sequence statistics and functional implications in gene expression. Nucleic Acids Res, 1987, 15:7155–74

16. Jaganathan K, Kyriazopoulou Panagiotopoulou S, McRae JF, Darbandi SF, Knowles D, Li YI, Kosmicki JA, Arbelaez J, Cui W, Schwartz GB, Chow ED, Kanterakis E, Gao H, Kia A, Batzoglou S, Sanders SJ, Farh KK-H. Predicting Splicing from Primary Sequence with Deep Learning. Cell, 2019, 176:535–48.e24

17. Dobin A, Davis CA, Schlesinger F, Drenkow J, Zaleski C, Jha S, Batut P, Chaisson M, Gingeras TR. STAR: ultrafast universal RNA-seq aligner. Bioinformatics, 2012, 29:15–21

18. Smith T, Heger A, Sudbery I. UMI-tools: modeling sequencing errors in Unique Molecular Identifiers to improve quantification accuracy. Genome Res, 2017, 27:491–9

19. Katz Y, Wang ET, Airoldi EM, Burge CB. Analysis and design of RNA sequencing experiments for identifying isoform regulation. Nat Methods, 2010, 7:1009–15

20. Schafer S, Miao K, Benson CC, Heinig M, Cook SA, Hubner N. Alternative splicing signatures in RNA-seq data: Percent spliced in (PSI). Curr Protoc Hum Genet, 2015, 87:11.16.1–11.16.14

21. Martincorena I, Roshan A, Gerstung M, Ellis P, Van Loo P, McLaren S, Wedge DC, Fullam A, Alexandrov LB, Tubio JM, Stebbings L, Menzies A, Widaa S, Stratton MR, Jones PH, Campbell PJ. Tumor evolution. High burden and pervasive positive selection of somatic mutations in normal human skin. Science, 2015, 348:880–6

22. Horton C, Cass A, Conner BR, Hoang L, Zimmermann H, Abualkheir N, Burks D, Qian D, Molparia B, Vuong H, LaDuca H, Grzybowski J, Durda K, Pilarski R, Profato J, Clayback K, Mahoney M, Schroeder C, Torres-Martinez W, Elliott A, Chao EC, Karam R. Mutational and splicing landscape in a cohort of 43,000 patients tested for hereditary cancer. NPJ Genom Med, 2022, 7:49

23. Schröder S, Wieland B, Ohlenbusch A, Yigit G, Altmüller J, Boltshauser E, Dörk T, Brockmann K. Evidence of Pathogenicity for the Leaky Splice Variant c.1066-6T>G in ATM in a Patient with Variant Ataxia Telangiectasia. Abstracts of the 46th Annual Meeting of the Society for Neuropediatrics, 2021. https://doi.org/10.1055/s-0041-1739584

24. Bergsma AJ, Kroos M, Hoogeveen-Westerveld M, Halley D, van der Ploeg AT, Pijnappel WW. Identification and characterization of aberrant GAA pre-mRNA splicing in pompe disease using a generic approach. Hum Mutat, 2015, 36:57–68

