## Supplementary material for "A systematic method for detecting abnormal mRNA splicing and assessing its clinical impact in individuals undergoing genetic testing for hereditary cancer syndromes": Table S1

**Supplemental Table 1. Genes included in RNA analysis assay**

| **Gene** | **Reportable Transcript** | **Mechanism** | **% Junctions** | **Final Assay** | **Stated Limitations** |
| --- | --- | --- | --- | --- | --- |
| AIP | NM_003977.3 | LOF | 100 | Yes |  |
| ALK | NM_004304.4 | GOF | 0 | No |  |
| APC | NM_000038.5 | LOF | 100 | Yes |  |
| ATM | NM_000051.3 | LOF | 100 | Yes |  |
| AXIN2 | NM_004655.3 | LOF | 100 | Yes |  |
| BAP1 | NM_004656.3 | LOF | 100 | Yes |  |
| BARD1 | NM_000465.3 | LOF | 100 | Yes |  |
| BLM | NM_000057.3 | LOF | 100 | Yes |  |
| BMPR1A | NM_004329.2 | LOF | 100 | Yes |  |
| BRCA1 | NM_007294.3 | LOF | 100 | Yes |  |
| BRCA2 | NM_000059.3 | LOF | 100 | Yes |  |
| BRIP1 | NM_032043.2 | LOF | 100 | Yes |  |
| CASR | NM_000388.3 | LOF | 0 | No |  |
| CDC73 | NM_024529.4 | LOF | 100 | Yes |  |
| CDH1 | NM_004360.3 | LOF | >75 | Yes | Sensitivity involving exons 1-3 may be reduced. |
| CDK4 | NM_000075.3 | GOF | 100 | No |  |
| CDKN1B | NM_004064.4 | LOF | 100 | Yes |  |
| CDKN1C | NM_000076.2 | LOF | 100 | Yes |  |
| CDKN2A | NM_000077.4 | LOF | >40 | No |  |
| CDKN2A | NM_058195.3 | LOF | >40 | No |  |
| CEBPA | NM_004364.4 | LOF | No introns | No |  |
| CHEK2 | NM_007194.3 | LOF | 100 | Yes |  |
| CTNNA1 | NM_001903.3 | LOF | 100 | Yes |  |
| DICER1 | NM_177438.2 | LOF | 100 | Yes |  |
| DIS3L2 | NM_152383.4 | LOF | 100 | Yes |  |
| EGFR | NM_005228.3 | LOF | 0 | No |  |
| EPCAM | NM_002354.2 | LOF | >40 | No |  |
| FH | NM_000143.3 | LOF | 100 | Yes |  |
| FLCN | NM_144997.5 | LOF | 100 | Yes |  |
| GATA2 | NM_032638.4 | LOF | 100 | Yes |  |
| GPC3 | NM_004484.3 | LOF | 0 | No |  |
| GREM1 | NM_013372.6 | GOF | No Introns | No |  |
| HOXB13 | NM_006361.5 | unknown | 0 | No |  |
| HRAS | NM_005343.2 | GOF | 100 | No |  |
| KIT | NM_000222.2 | LOF | >75 | No |  |
| MAX | NM_002382.4 | LOF | 100 | Yes |  |
| MEN1 | NM_130799.2 | LOF | 100 | Yes |  |
| MET | NM_001127500.1 | unknown | 0 | No |  |
| MITF | NM_000248.3 | LOF | >75 | No | Single Site DNA test |
| MLH1 | NM_000249.3 | LOF | 100 | Yes |  |
| MSH2 | NM_000251.2 | LOF | 100 | Yes |  |
| MSH3 | NM_002439.4 | LOF | 100 | Yes |  |
| MSH6 | NM_000179.2 | LOF | 100 | Yes |  |
| MUTYH | NM_001128425.1 | LOF | >75 | Yes | Sensitivity involving exons 2-3 may be reduced. |
| NBN | NM_002485.4 | LOF | 100 | Yes |  |
| NF1 | NM_000267.3 | LOF | 100 | Yes |  |
| NF2 | NM_000268.3 | LOF | 100 | Yes |  |
| NTHL1 | NM_002528.6 | LOF | 100 | Yes |  |
| PALB2 | NM_024675.3 | LOF | 100 | Yes |  |
| PDGFRA | NM_006206.4 | unknown | <40 | No |  |
| PHOX2B | NM_003924.3 | LOF | 0 | No |  |
| PMS2 | NM_000535.5 | LOF | 100 | Yes | Sensitivity involving exons 11-15 may be reduced. |
| POLD1 | NM_002691.3 | LOF | 100 | Yes |  |
| POLE | NM_006231.3 | LOF | 100 | Yes |  |
| POT1 | NM_015450.2 | LOF | >75 | Yes |  |
| PRKAR1A | NM_002734.4 | LOF | 100 | Yes |  |
| PTCH1 | NM_000264.3 | LOF | 100 | Yes |  |
| PTEN | NM_000314.4 | LOF | 100 | Yes |  |
| RAD50 | NM_005732.3 | LOF | 100 | Yes |  |
| RAD51C | NM_058216.2 | LOF | 100 | Yes |  |
| RAD51D | NM_002878.3 | LOF | 100 | Yes |  |
| RB1 | NM_000321.2 | LOF | 100 | Yes |  |
| RECQL4 | NM_004260.3 | LOF | <40 | No |  |
| RET | NM_020975.4 | LOF | <40 | No |  |
| RUNX1 | NM_001754.4 | LOF | 100 | Yes |  |
| SDHA | NM_004168.3 | LOF | 100 | Yes |  |
| SDHAF2 | NM_017841.2 | LOF | 100 | Yes |  |
| SDHB | NM_003000.2 | LOF | 100 | Yes |  |
| SDHC | NM_003001.3 | LOF | 100 | Yes |  |
| SDHD | NM_003002.3 | LOF | 100 | Yes |  |
| SMAD4 | NM_005359.5 | LOF | 100 | Yes |  |
| SMARCA4 | NM_001128849.1 | LOF | 100 | Yes |  |
| SMARCB1 | NM_003073.3 | LOF | 100 | Yes |  |
| SMARCE1 | NM_003079.4 | LOF | 100 | Yes |  |
| STK11 | NM_000455.4 | LOF | 100 | Yes |  |
| SUFU | NM_016169.3 | LOF | 100 | Yes |  |
| TERC | NR_001566.1 | LOF | No Intron | No |  |
| TERT | NM_198253.2 | LOF | 0 | No |  |
| TMEM127 | NM_017849.3 | LOF | 100 | Yes |  |
| TP53 | NM_000546.5 | LOF | 100 | Yes |  |
| TSC1 | NM_000368.4 | LOF | 100 | Yes |  |
| TSC2 | NM_000548.3 | LOF | >75 | Yes | Sensitivity involving exons 25-27 may be reduced. |
| VHL | NM_000551.3 | LOF | 100 | Yes |  |
| WRN | NM_000553.4 | LOF | 100 | Yes |  |
| WT1 | NM_024426.4 | LOF | 0 | No |  |
