## Supplementary material for "A systematic method for detecting abnormal mRNA splicing and assessing its clinical impact in individuals undergoing genetic testing for hereditary cancer syndromes": Table S2

**Supplemental Table 2: Sherloc evidence for interpretation of RNA effects**

| **Sherloc evidence description** | **P score** | **B score** |
| --- | --- | --- |
| Altered splicing (in vivo), NMD expected [LoF] | 3 |  |
| Altered splicing (in vivo), NMD expected [not-LoF] | 2 |  |
| Altered splicing (in vitro or in vivo), out-of-frame, may escape NMD [LoF] | 3 |  |
| Altered splicing (in vitro or in vivo), out-of-frame, may escape NMD [not-LoF] | 1 |  |
| Altered splicing (in vitro or in vivo), initiator skipping [LoF] | 2.5 |  |
| Altered splicing (in vitro or in vivo), initiator skipping [not-LoF] | 2 |  |
| Altered splicing (in vitro or in vivo), in-frame exon-skipping [LoF] | 3 |  |
| Altered splicing (in vitro or in vivo), in-frame exon-skipping [not-LoF] | 1 |  |
| Altered splicing (in vitro or in vivo), small in-frame loss (loss of one or more codons) | 1 |  |
| Altered splicing (in vitro or in vivo), in-frame insertion (gain of one or more codons) | 1 |  |
| Altered splicing (in vitro or in vivo), complex | 3 |  |
| No significantly altered splicing observed |  | 1 |
| AddOn: One or more observed mRNA has previously been described as a naturally-occurring isoform |  | 0 |
| In vitro splicing: Evidence for resulting in NMD+ variant | 1 |  |
| In vitro splicing: Weak evidence for no impact |  | 0 |
